# A Spatial Omnibus Test (SPOT) for Spatial Proteomic Data

**DOI:** 10.1101/2024.03.08.584117

**Authors:** Sarah Samorodnitsky, Katie Campbell, Antoni Ribas, Michael C. Wu

## Abstract

Spatial proteomics can reveal the spatial organization of immune cells in the tumor immune microenvironment. Relating measures of spatial clustering, such as Ripley’s K or Besag’s L, to patient outcomes may offer important clinical insights. However, these measures require pre-specifying a radius in which to quantify clustering, yet no consensus exists on the optimal radius which may be context-specific. We propose a SPatial Omnibus Test (SPOT) which conducts this analysis across a range of candidate radii. At each radius, SPOT evaluates the association between the spatial summary and outcome, adjusting for confounders. SPOT then aggregates results across radii using the Cauchy combination test, yielding an omnibus p-value characterizing the overall degree of association. Using simulations, we verify that the type I error rate is controlled and show SPOT can be more powerful than alternatives. We also apply SPOT to an ovarian cancer study. An R package and tutorial is provided at https://github.com/sarahsamorodnitsky/SPOT.

## 1 Introduction

Popular proteomic imaging platforms, such as multiplexed ion beam imaging (MIBI), multiplexed immunohistochemistry (mIHC), and imaging mass cytometry (IMC), can be used to examine the spatial distribution of cells in tissue [1]. These platforms record the spatial location and protein marker expression levels – used to identify types and activities – of cells within the tissue. This permits study of the tumor immune microenvironment (TIME), the landscape of immune cells within a tumor, at the single-cell and spatial level [2], which has been shown to be associated with clinical outcomes, such as survival [3, 4, 5], response to treatment [3, 6], and disease recurrence [4].

A common analytical approach to studying the spatial distribution of cells leverages the homogeneous point process model which characterizes whether cells are randomly distributed around the image (i.e. exhibiting *complete spatial randomness* or CSR), clustered, or dispersed/repulsed [1]. This model allows us to test the assumption that cells (which may be labeled by their cell type, e.g. CD8 T cell) are distributed under CSR and characterize the spatial organization of cells, offering potentially useful insights. To do so, we can summarize the spatial organization of cells within a radius *t* using spatial summary statistics like Ripley’s K [7], Besag’s L [8], and Marcon’s M [9]. Bivariate generalizations can be used to characterize the spatial colocalization of two cell types [10]. Bivariate Ripley’s K, Besag’s L, and Marcon’s M quantify the degree of co-occurrence between two cell types and test whether two cell types are clustered together, dispersed, or randomly distributed under CSR [1].

A challenge with these spatial measures is the choice of radius, *t*, for characterizing proximal relationships. This could be guided by clinical knowledge [11, 12], but there is no consensus or guideline across applications and hypotheses. Fixing the radius at one value may neglect clinically relevant spatial patterns observed at smaller or larger values of *t*. Prior work has considered a functional analytic approach [13, 14] in which spatial summary measures evaluated at multiple radii are treated as functional covariates in an outcome model. However, this requires several tuning parameters, may be computationally intensive, and may make interpretations difficult for clinicians. Alternatively, spicyR [15] considers a range of radii and produces an overall summary measure, termed a colocalization score, by calculating the area between the estimated spatial statistic and the theoretical value under CSR across radii, which is treated as the outcome or response variable. While this accommodates multiple radii, it does not easily accommodate censored clinical outcomes, like overall survival. Dayao et al. [16] treat *t* as a tuning parameter and select several values to use based on the concordance index. This still requires choosing a radius at which to interpret the results and may be challenging to synthesize an interpretation across multiple radii. This afflicts both univariate and bivariate colocalization analyses.

To address the challenge in selecting a radius, we propose an alternative approach, the SPatial Omnibus Test (SPOT). SPOT involves three steps: first, the user provides a series of radii at which to calculate a spatial summary statistic (e.g. Ripley’s K, Besag’s L); then, the association between the spatial summary and a clinical phenotype, like survival or treatment response, is tested at each radius using an appropriate model (e.g. Cox proportional hazards for a survival outcome with the spatial summary as a covariate) which results in a p-value describing association for each radius; finally, the p-values across radii are combined using the Cauchy combination test [17]. This yields a single “omnibus” p-value characterizing the overall strength of association between spatial organization of cells and patient outcomes. The power for detecting this association depends heavily on the user’s choice of radius, such that choosing a poor radius results in low power. On the other hand, considering multiple radii and choosing the radius resulting in greatest statistical significance leads to severe false positives. Thus, the advantages of SPOT are that, as an omnibus test, it precludes need to choose a radius *a priori*, protects the false positive rate, and maintains high power.

The rest of this article is organized as follows. In Section 2, we introduce the SPOT method. In Section 3, we use SPOT to evaluate the relationship between immune cell clustering in ovarian cancer. In Section 4, we compare the power of SPOT against testing at individual fixed radii and the type I error rate and power of SPOT compared to existing methods. We conclude with a brief discussion in Section 5.

## 2 Methods

Suppose we have *M* tumor samples, which we index by *m* = 1, …, *M*. For each sample, we may have *R*_*m*_ *≥* 1 regions-of-interest (ROIs) or images which show the spatial location of the detected cells and their phenotypes within a specific sub-region of the tumor sample. We index ROIs within a tumor sample using *r* = 1, …, *R*_*m*_. For sample *m* and ROI *r*, we assume there are *n*_*mr*_ detected cells, irrespective of cell-type label. We index these cells using *i* and *j*, i.e. *i, j* = 1, …, *n*_*mr*_. For cell types *a* and *b*, we assume there are 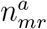 and 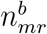 of each in ROI *r* in sample *m*, respectively. Let **t** = (*t*_1_, …, *t*_*P*_) define a vector of radii where *t*_1_ = 0.

Our goal is to quantify the strength of association between the spatial organization of cells in tumors with a clinical phenotype. To do so, we follow a three-step procedure:

1. Select a series of radii, **t**, and a spatial summary measure to characterize the spatial distribution of cells. Evaluate the spatial summary measure at each radius *t*_*p*_ for each sample *m* and ROI *r*.
2. Test the association between the spatial summary at radius *t*_*p*_ on the clinical outcome after adjustment for clinical covariates, like age or sex, using the appropriate outcome model (e.g. Cox proportional hazards model or logistic regression). Repeat this process for *t*_1_, …, *t*_*P*_.
3. Combine the resulting *P* p-values using the Cauchy combination test. This provides an omnibus p-value describing the association between the spatial summary across radii and the clinical outcome.

Next, we will describe each step to our approach in more detail. We frame our description of the SPOT method using Besag’s L as a spatial summary measure and survival as the clinical outcome. However, this framework is general and can accommodate many spatial summary statistics and patient outcomes best suited to the scientific question at hand.

### 2.1 Spatial Summary Measures

We treat the cell locations in each ROI as a spatial point pattern, which is a realization of a point process [18]. This point process may be unmarked, meaning we disregard or do not possess additional cell-level information. This information could include functional or phenotypic marker expression, e.g. the expression of cytokeratin (CK), or categorical cell type labels, e.g. tumor cell. We focus our discussion on categorical marks, but this framework accommodates continuous marks, as well. Note that for our purposes each point in a point pattern is a cell so we will use “point” and “cell” interchangeably.

We can describe the spatial organization of the point pattern within an ROI using second-order spatial summary statistics, which characterize the expected number of points within a radius *t*. Ripley’s K and Besag’s L are two related examples of second-order spatial summaries, which we will leverage to characterize the spatial organization of cells across tumor samples. These summary statistics are functions which take in the *n*-dimensional cell locations and output a scalar value. This value indicates how closely the point pattern adheres to the assumption of *complete spatial randomness* or CSR, when the point (cell) locations are independent of each other. If the point pattern deviates from CSR, it may exhibit clustering or dispersion. In this case, Ripley’s K and Besag’s L quantify the degree of clustering or dispersion within the pattern.

We first define Ripley’s K, denoted by *K*(*t*), using the notation given in Dixon (2002) [19]. Under complete spatial randomness and for a given radius *t, K*(*t*) = *πt*^2^. For a homogeneous point process, in which points or cells are equally likely to arise anywhere within the ROI, we can estimate *K*(*t*) for ROI *r* and sample *m* by:

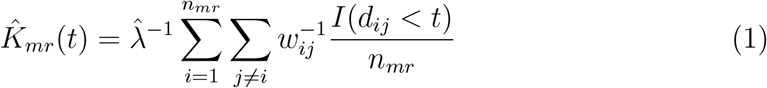

where 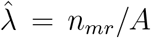, *A* is the area of the study region, *d*_*ij*_ is the distance between points *i* and *j, w*_*ij*_ is an edge correction in cases when the circle of radius *t* crosses the edge of the study region, and *I*(*d*_*ij*_ *< t*) = 1 if *d*_*ij*_ *< t* and 0 otherwise. In our data application (Section 3), we use Ripley’s isotropic edge correction [10] to correct for edge effects. The estimate given in Equation 1 treats the cells as unlabeled. For a labeled subset of cells of type *a* (e.g., CD8 T cells only), we can adjust 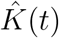 accordingly:

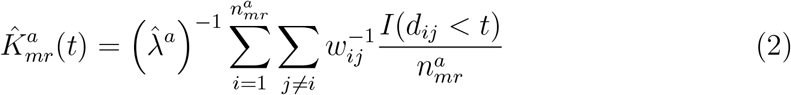

where 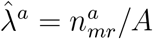

Besag’s L, denoted by *L*(*t*), is closely related to Ripley’s K. Under complete spatial randomness, 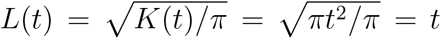 Our empirical estimate of *L*(*t*) for sample *m* and ROI *r* is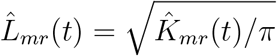. For cell type *a*, our estimate would be 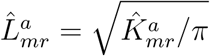.

Ripley’s K and Besag’s L have bivariate generalizations to characterize the expected number of colocations between two point (cell) types. The bivariate generalization of Ripley’s K is:

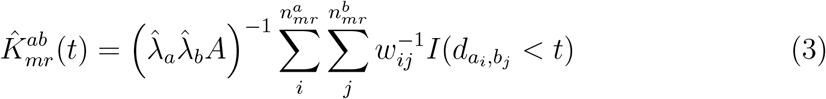

where 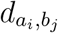 is the distance between the *i*th point of type *a* and the *j*th point of type *b*. The bivariate generalization of Besag’s L would then be 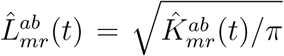 For the remainder of this article, we focus on Besag’s L as a measure of the spatial distribution of cells.

To select **t**, we use the recommended default given in the `spatstat R` package referred to as “Ripley’s rule of thumb.” This suggests using a range of radii between 0 and 0.25 times the shortest side of the image [20]. In practice, some spatial summaries, particularly for small values of *t*_*p*_, will be 0. We suggest only including radii for which at least 20% of images have a non-zero spatial summary value. We followed this reasoning in our data application in Section 3.

### 2.2 Outcome Model

In our discussion, we focus on a survival outcome but this framework accommodates other outcome models, as well. We use a Cox proportional hazards model to examine the effect spatial summary of cells or colocalization of cell types as measured by Besag’s L on the log-hazard of an event. We adjust for additional clinical covariates that may confound this relationship. For a given radius, *t*_*p*_, define 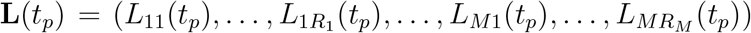 as a vector of Besag’s L evaluated at radius *t*_*p*_ across samples and ROIs. If *R*_*m*_ = 1 for all *m* = 1, …, *M*, **L**(*t*_*p*_) = (*L*_1_(*t*_*p*_), …, *L*_*M*_ (*t*_*p*_)). If *R*_*m*_ *>* 1 for some *m*, we average *L*_*mr*_(*t*_*p*_) within each sample *m*, i.e. 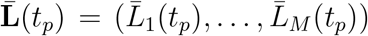 where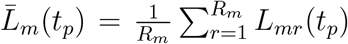. In addition to **L**(*t*_*p*_) we may also have *B* clinical covariates contained in a design matrix **X** : *M × B*. Our Cox proportional hazards model can be specified as:

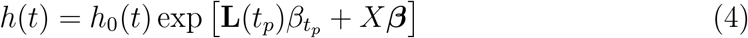

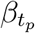 reflects the effect of the spatial configuration of cells or cell types within the ROIs on the log-hazard for the event-of-interest. We store the p-value derived from a standard Wald test of *H*_0_ : 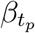 = 0. For each radius, *t*_1_, …, *t*_*p*_, we obtain *P* p-values, *p*_1_, …, *p*_*P*_.

### 2.3 Omnibus Test

We have a series of *P* p-values, *p*_1_, …, *p*_*P*_, describing the significance of the effect of **L**(*t*_*p*_) on the log-hazard for the event. Our goal is to obtain a summary p-value based on *p*_1_, …, *p*_*P*_ that describes the overall significance of the effect of Besag’s L across radii *t*_1_, …, *t*_*P*_ on the log-hazard.

To obtain an overall p-value based on *p*_1_, …, *p*_*P*_, we use the Cauchy combination test [17]. The Cauchy combination test was developed to combine multiple, potentially dependent, p-values into one summary value. For p-values *p*_1_, …, *p*_*P*_, the Cauchy combination test statistic is defined as:

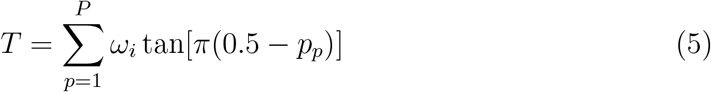

where *ω*_*p*_ represents a weight, which we fix *ω*_*p*_ = 1*/P* for all *p*. Under the null, each tan[*π*(0.5 *− p*_*p*_)] follows a standard Cauchy distribution.

## 3 Data Application

We used SPOT to analyze images from 128 patients with high-grade serous caricoma (HGSOC) [21]. The dataset was generated using the Vectra-Polaris multi-spectral immunohistochemistry (IHC) platform. We retrieved the dataset from http://juliawrobel.com/MI_tutorial/MI_Data.html but this data can also be downloaded from the VectraPolarisData R package [22]. The detected immune cell types were B cells (cells positive for the CD19 marker or CD19+), macrophages (CD68+), CD4 T cells (CD3+, CD8-), and CD8 T cells (CD3+, CD8+). Tumor cells were identified if cells were positive for the CK marker (CK+). The cell locations were categorized as being within the tumor region or the stroma region of each ROI. For our analysis, we only considered cells located within the tumor region.

We tested for associations between the spatial configuration of each immune cell type (B cells, macrophages, CD4 T cells, CD8 T cells) with overall survival, adjusting for patient age at diagnosis. We also tested for an association between spatial colocalization of each pair of immune cells with overall survival, adjusting for patient age. Within both analyses, we adjusted the p-values for multiple testing using an FDR adjustment. Since the dimensions of the images varied, we choose a range of radii between 0 and 0.25 times the smallest image width across the images which was 0.25 *∗* 1009.6 = 252.4. We considered 100 different radii between 0 and 252.4.

We found that the clustering of each immune cell type individually was not associated with overall survival. The FDRs for each cell type were *p* = 0.261 for macrophages, *p* = 0.486 for B cells, *p* = 0.477 for CD8 T cells, and *p* = 0.936 for CD4 T cells. Within a short window (*t*_*p*_ *∈* (2.55, 12.75)), the association between macrophage clustering and overall survival appeared significant but dissipated as the radius increased.

We then tested the association between the colocalization of pairs of immune cells and overall survival. We found that the colocalization of CD4 T cells and macrophages was significantly associated with overall survival (FDR = 0.0286). We also found that the colocalization of macrophages and B cells was significantly associated with overall survival (FDR = 0.0286) (Table 1, Figure 1). These results align with Steinhart et al.’s [21] analysis of this dataset, which revealed that proximity between macrophages and B cells and between macrophages and CD4 T cells was associated with overall survival.

**Table 1:** Association between the colocalization of each immune cell type pair and overall survival in ovarian cancer.

| Cell Type 1 | Cell Type 2 | SPOT P-Value | FDR |
| --- | --- | --- | --- |
| CD4 T cell | Macrophage | 0.0071 | 0.0286 |
| Macrophage | B cell | 0.0095 | 0.0286 |
| CD4 T cell | CD8 T cell | 0.1887 | 0.3774 |
| Macrophage | CD8 T cell | 0.3959 | 0.5938 |
| CD8 T cell | B cell | 0.9191 | 0.9216 |
| CD4 T cell | B cell | 0.9216 | 0.9216 |

**Figure 1:**
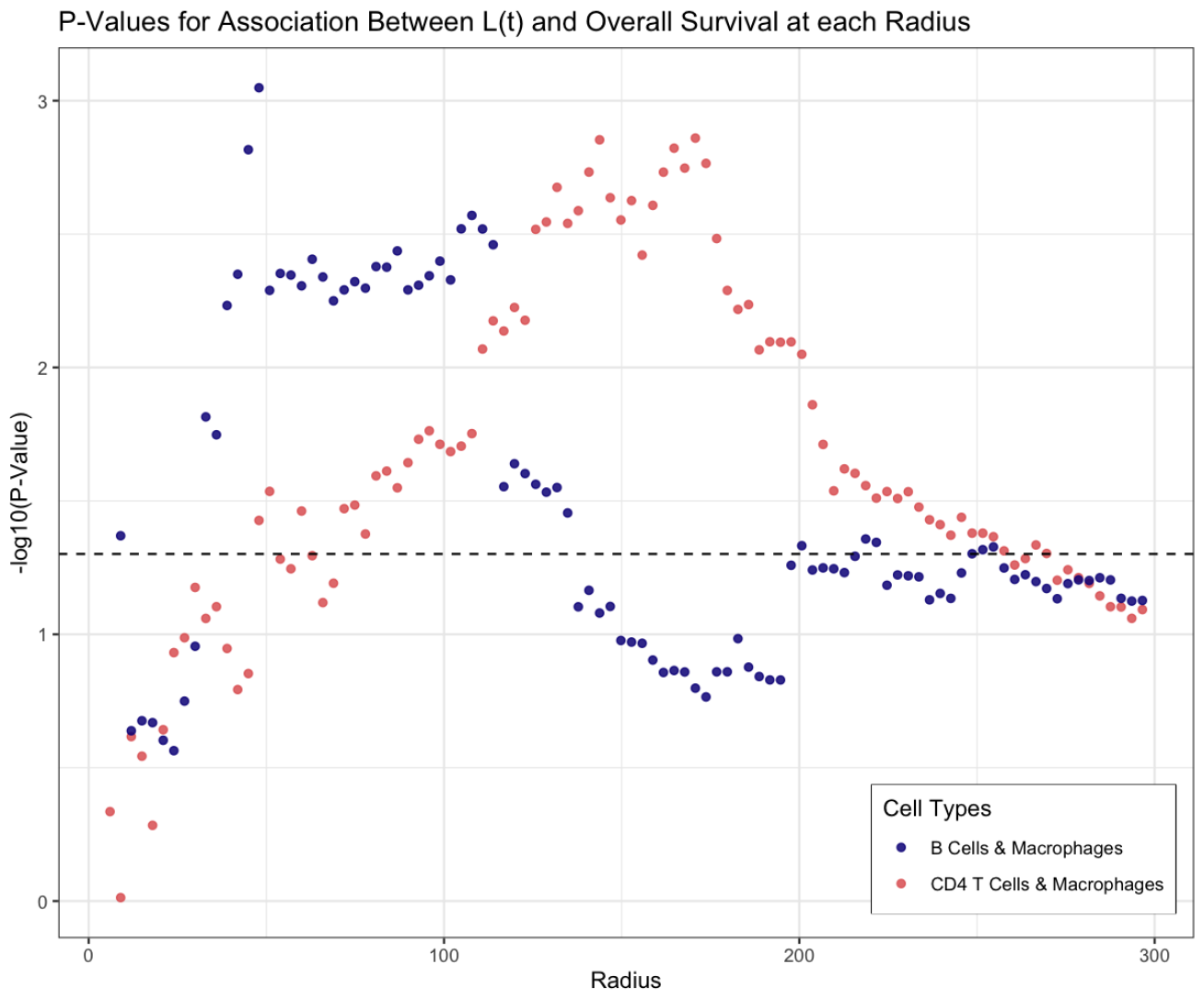
The relationship between the radius, *t*, and the p-value for the effect of CD4 T cell-macrophage colocalization and macrophage-B cell colocalization as measured by *L*(*t*) on overall survival.

## 4 Simulation Study

### 4.1 Power at each Fixed Radius

We first evaluated the power of testing the association between a spatial summary at a series of individual radii with survival against SPOT’s power. For this simulation, we used Besag’s L as our spatial summary measure. We generated images for *M* = 100 samples. To generate a survival outcome, we simulated the event times for the first 50 samples from Exponential(*λ* = log(2)*/*12) (the low-survival group) and the event times for the last 50 samples from Exponential(*λ* = 0.4 log(2)*/*12) (the high-survival group) where *λ* represents the rate or hazard parameter. We randomly censored 10-20% of event times in each group. Each image was generated to have dimension 1000 *×* 1000. We generated between 50 and 100 cells in each image. We randomly labeled each cell as one of two cell types so that there were approximately equal numbers of each type. We generated the cell locations for each image under two conditions: (1) exhibiting complete spatial randomness or (2) exhibiting spatial clustering. To generate an image under CSR, we simulated the (*x, y*) coordinates for each cell from a uniform distribution, i.e. *x, y ∼* Uniform(0, 1000). To generate a clustered image, we first simulated mean *x* and *y* locations from a uniform distribution, i.e. *µ*_*x*_, *µ*_*y*_ *∼* Uniform(100, 900), and then simulated the (*x, y*) coordinates for each cell from a multivariate-normal distribution: (*x, y*) *∼* Multivariate-Normal((*µ*_*x*_, *µ*_*y*_)^*T*^, **Σ**) where 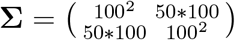

To estimate power, we simulated images for the higher-survival group to be uniform and images for the low-survival group to be clustered. We considered the power at each radius value from 0 to 250. We ran the simulation for 1000 replications and calculated power using the proportion of simulation replications the p-values were below a significance level of 0.05.

The results are shown in Figure 2. As the radius increases, we observe a gain in power across the radii before a slight descent. The maximum individual power was 0.853 at a radius of 189 whereas SPOT’s power was 0.867. This illustrates the variability in power observed at each radius and the challenge of choosing the “best” radius at which to relate the spatial distribution of cells with clinical outcomes.

**Figure 2:**
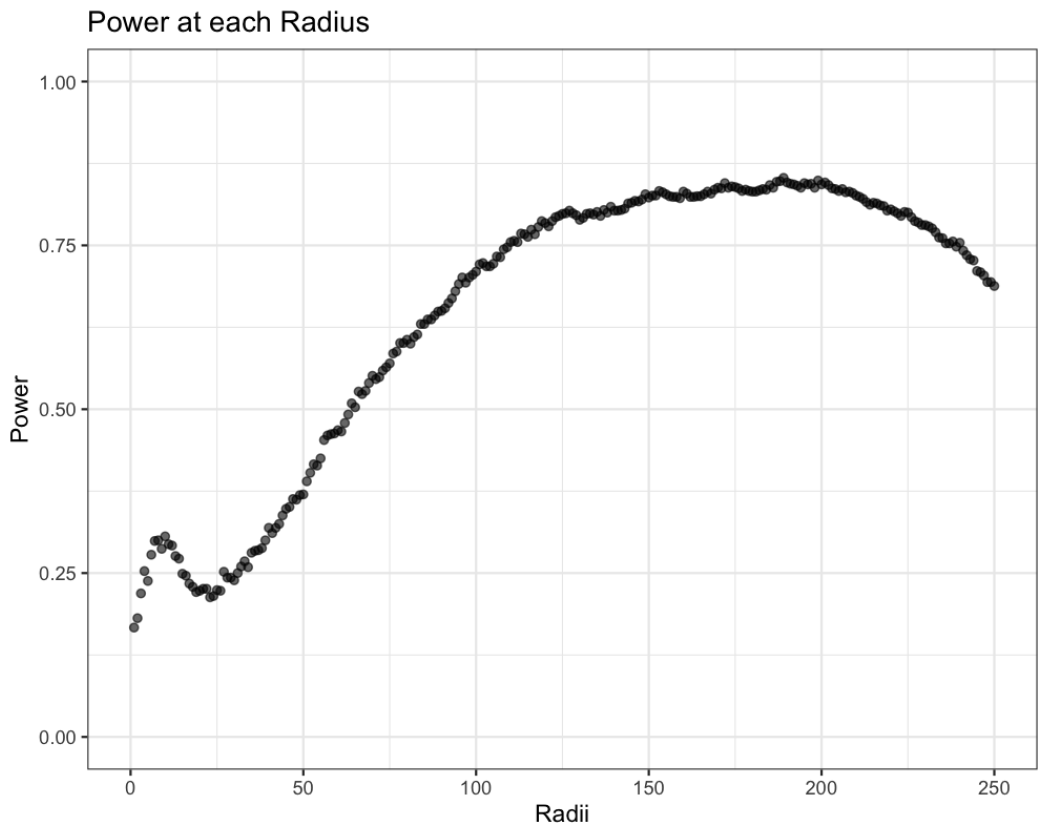
The power of testing the association between Besag’s L at each radius and survival.

### 4.2 Methods Comparison

We then compared SPOT to existing methods. We considered SPF [13] and FunSpace [14] as alternative approaches. These methods allow for a range of radii and treat the spatial summary measure evaluated at each radius as a functional covariate. We also compared these methods against a “naive” approach in which we select the “best” and “worst” radius at which to calculate a spatial summary. For SPOT and the naive approach, we use Besag’s L as our spatial summary statistic. We compared the approaches in terms of their type I error rate and power across several conditions.

Throughout the simulation, we varied the:

1. Outcome type (survival or binary)
2. Number of cell types (one or two)
3. Number of images per sample (one or multiple)

Here we focus on the results for a survival outcome and a single image per sample. In Section 1.1 of the Supplementary Materials, we consider simulating multiple images and in Section 1.2 we consider a binary outcome. Under each condition, we generated images of dimension 1000 *×* 1000 for *M* = 100 samples and generated a survival outcome in an analogous manner as described in Section 4.1.

We fit SPF and FunSpace in the following manner. To fit SPF with one cell type, we used the pair correlation function [10] as the spatial summary measure for an unmarked point process. For two cell types, we used the mark connection function as described in Vu et al. [13]. When we generated multiple images per sample, we averaged the pair correlation function or mark connection function output across ROIs within a sample. We also extended SPF to allow for a binary outcome. We fit FunSpace only for conditions with two cell types and one image per sample. We also extended FunSpace to allow for a binary outcome.

To estimate the type I error rate, images were randomly generated to be uniform or clustered. To estimate power under conditions where each sample had only one image, we simulated images for the high-survival group to be uniform and images for the low-survival group to be clustered. To estimate power under conditions in which there were multiple images per sample, we simulated all images to be uniform for the high-survival group. For the low-survival group we randomly chose some images to be clustered and some to be uniform. The probability of an image being generated as clustered in this group was 0.75. We calculated type I error and power using the proportion of simulation replications the p-values were below a significance level of 0.05.

We ran each simulation condition for 1000 replications. The results for one image per sample are shown in Table 2. With both one and two cell types, SPOT exhibits the lowest type I error rate (one cell type: 0.057 for SPOT vs. 0.065 for SPF; two cell types: 0.054 vs. 0.057 for SPF and 0.059 for FunSpace). At the “best” radius, the power for one cell type was 0.992 and 0.991 for two cell types, though the type I error rate was 0.456 for one cell type and 0.483 for two cell types. At the “worst” radius, the power for one cell type was 0 and 0.001 for two cell types, though the type I error rate was 0.006 for one cell type and 0.005 for two cell types. This illustrates that SPOT provides a middle-ground between choosing the “best” or ideal radius in every condition, at which the type I error rate is high, and the “worst” or most-conservative radius, at which power is very low.

**Table 2:** Type I error rates and power with a survival outcome and one image per sample.

| Model | 1 Cell Type |  | 2 Cell Types |  |
| --- | --- | --- | --- | --- |
|  | Type I Error Rate | Power | Type I Error Rate | Power |
| SPOT | 0.057 | 0.925 | 0.054 | 0.867 |
| SPF | 0.065 | 0.951 | 0.057 | 0.305 |
| FunSpace |  |  | 0.059 | 0.764 |
| Choose ‘Best’ Radius | 0.456 | 0.992 | 0.483 | 0.991 |
| Choose ‘Worst’ Radius | 0.000 | 0.006 | 0.001 | 0.005 |

## 5 Discussion

We proposed a SPatial Omnibus Test (SPOT) for the association between spatial summary measures of cell organization in the TIME with clinical outcomes, like survival or treatment response. These summary measures typically require the user to select a radius at which to characterize spatial organization of cells. However, there is no rule-of-thumb or guideline for making this selection across applications. SPOT provides a straightforward framework for relating the cellular architecture (or cell type colocalization) with outcomes across multiple radii. We found via simulation that SPOT provides a reasonable middle-ground between choosing the “best” radius, which is difficult to know *a priori* and may lead to false positive discoveries, and the “worst” radius, which offers very little power. We also applied SPOT to an ovarian cancer dataset, which corroborated the prognostic importance of CD4 T cell and macrophage colocalization, as well as macrophage and B cell colocalization [21].

The advantage of the SPOT framework is that it is adaptable to the application and hypotheses of practitioners. For example, one could consider any measure of spatial cellular configuration, such as the mark connection function used by Vu et al. [13] or estimate the spatial intensity of cells and consider the Jensen-Shannon distance between density estimates as done in Masotti et al. [12]. One could consider spatial summary measures that accommodate inhomogeneity and could implement any outcome model depending on the clinical outcome-of-interest. Further, SPOT could easily be parallelized across radii to improve computation speed.

We used Ripley’s rule-of-thumb for determining radii ranges. Our data analyses suggest that this range is reasonable, but one could expand beyond this. Consideration of additional radii can boost power if useful radii are included, but consideration of poor choices can lead to reduced power. Thus, if one has prior contextual knowledge, applying SPOT to a more targeted, reduced set of radii may offer improvement.

We emphasize that we focused on Besag’s L, but SPOT can be applied to any relevant and valid summary statistic for convenience. The framework remains the same, simply substituting the choice of metric. In principle, we could further extend SPOT to simultaneous consideration of multiple metrics, as well as radii, but this remains a potential topic for further investigation. Finally, we did not address the issue of holes or gaps in the image that may arise in tumor resections using spatial proteomics imaging platforms. The challenge of gaps in the image is that it violates the assumption of homogeneity among the points in a spatial point pattern. One approach to address this is to incorporate a simulation envelope in which the cell type labels are permuted [23, 24]. This approach could be incorporated into SPOT, though the computational cost remains high. As further approaches are developed in this area, we anticipate their potential incorporation into our proposed framework and the continued growth of our method.

## 7 Acknowledgments

Funding: This work was supported in part by NIH Grant U10 CA180819 and The Hope Foundation for Cancer Research.

