## Supplementary Materials for "A Spatial Omnibus Test (SPOT) for Spatial Proteomic Data"

#### 1 Simulation Studies

##### 1.1 Multiple Images

We now describe our simulation results when we considered a survival outcome and multiple images per sample. Images were generated as discussed in Section 4 of the main manuscript. We randomly chose the number of images to simulate for each sample between one and three.

The results are shown in Table 1. With one cell type, SPOT exhibited slightly elevated type I error rates (0.065), though lower than SPF (0.074). This may be

| Model | 1 Cell Type | | 2 Cell Types | |
|---|---|---|---|---|
| | **Type I Error Rate** | **Power** | **Type I Error Rate** | **Power** |
| SPOT | 0.065 | 0.921 | 0.053 | 0.872 |
| SPF | 0.074 | 0.936 | 0.055 | 0.148 |
| Choose 'Best' Radius | 0.471 | 0.991 | 0.492 | 0.985 |
| Choose 'Worst' Radius | 0.000 | 0.003 | 0.000 | 0.008 |

Table 1: Type I error rates and power with a survival outcome and multiple images per sample

because the Cauchy combination test better controls type I error at lower significance levels. With two cell types, SPOT controlled type I error around 0.053 and exhibited much higher power than SPF (0.872 vs. 0.148). As discussed in the main manuscript, choosing the "best" radius yielded inflated type I error rates and the "worst" offered very little power. Similar to our results for a single image per sample, SPOT offers a balance between the "best" and "worst" radius in providing nominal type I error rates and adequate power.

#### 1.2 Binary Outcome

We consider the performance of SPOT against existing methods for relating spatial summary measures to a binary outcome with both a single image and multiple images per sample. We compared SPOT to SPF and FunSpace, as considered in the main manuscript, as well as SpaceANOVA [1]. SPF and FunSpace were not originally developed for a binary outcome. We extended these methods to leverage a functional logistic regression model. SpaceANOVA is another functional data analytic approach that treats a spatial summary measure evaluated across radii as a functional outcome.

Using a functional ANOVA model, SpaceANOVA incorporates a random effect in cases when there may be more than one image per sample. We fit SpaceANOVA only for conditions with two cell types and a binary outcome. We fixed the radii considered in SPF and SpaceANOVA at the same values considered for SPOT, which were from 0 to 250 by increments of 1.

The images were generated as described in the main manuscript. We assigned the first 50 samples to have an outcome of 1 (the "1-group") and the remaining 50 to have an outcome of 0 (the "0-group"). To assess type I error, we randomly simulated images as either uniform or clustered. To estimate power when each sample had only one image, we simulated images for the 1-group to be uniform and images for 0-group to be clustered. To estimate power when there may be more than one image per sample, all images in the 1-group were simulated to be uniform. For the 0-group we randomly chose some images to be clustered and some to be uniform.

The results for one image per sample are shown in Table 2. For both one and two cell types, SPOT controls type I error and offers power between 0.89 and 0.96, though SpaceANOVA provides the highest power.

| | 1 Cell Type | | 2 Cell Types | |
|---|---|---|---|---|
| **Model** | **Type I Error Rate** | **Power** | **Type I Error Rate** | **Power** |
| SPOT | 0.042 | 0.960 | 0.054 | 0.889 |
| SPF | 0.024 | 0.864 | 0.021 | 0.161 |
| FunSpace | | | 0.064 | 0.790 |
| SpaceANOVA | | | 0.062 | 0.999 |
| Choose 'Best' Radius | 0.396 | 0.999 | 0.465 | 0.991 |
| Choose 'Worst' Radius | 0.000 | 0.007 | 0.000 | 0.001 |

Table 2: Type I error rates and power for a binary outcome and one image per sample.

The results for multiple images per sample are shown in Table 3. SPOT controls type I error and provides adequate power. SpaceANOVA excels at controlling type I error and power under this condition.

| Model | 1 Cell Type | | 2 Cell Types | |
|---|---|---|---|---|
| | **Type I Error Rate** | **Power** | **Type I Error Rate** | **Power** |
| SPOT | 0.037 | 0.940 | 0.046 | 0.856 |
| SPF | 0.022 | 0.825 | 0.018 | 0.071 |
| SpaceANOVA | | | 0.048 | 1.000 |
| Choose 'Best' Radius | 0.426 | 1.000 | 0.472 | 0.987 |
| Choose 'Worst' Radius | 0.000 | 0.004 | 0.000 | 0.003 |

Table 3: Type I error rates and power for a binary outcome and multiple images per sample.

### References

[1] Souvik Seal, Brian Neelon, Peggi Angel, Elizabeth C O'Quinn, Elizabeth Hill, Thao Vu, Debashis Ghosh, Anand Mehta, Kristin Wallace, and Alexander V Alekseyenko. Spaceanova: Spatial co-occurrence analysis of cell types in multiplex imaging data using point process and functional anova. *bioRxiv*, 2023.
